# Active deep learning reduces annotation burden in automatic cell segmentation

**DOI:** 10.1101/211060

**Authors:** Aritra Chowdhury, Sujoy K. Biswas, Simone Bianco

## Abstract

The relationship between cellular architecture and cellular state and function is apparent, but not yet completely understood. Precise characterization of cellular state is important in many fields, from pathology to synthetic biology. High-content high-throughput microscopy is now more than ever accessible to researchers. This allows for collection of large amount of cellular images. Naturally, the analysis of this data cannot be left to manual investigation and needs to resort to the use of efficient computing algorithms for cellular detection, segmentation, and tracking. Annotation is required for building high quality algorithms. Medical professionals and researchers spend a lot of effort and time in annotating cells. This task has proved to be very repetitive and time consuming. The experts’ time is valuable and should be used effectively. Our hypothesis is that active deep learning will help to share some of the burden that researchers face in their everyday work. In this paper, we focus specifically on the problem of cellular segmentation.

We approach the segmentation task using a classification framework. Each pixel in the image is classified based on whether the patch around it resides on the interior, boundary or exterior of the cell. Deep convolutional neural networks (CNN) are used to perform the classification task. Active learning is the method used to reduce the annotation burden. Uncertainty sampling, a popular active learning framework is used in conjunction with CNN to segment the cells in the image. Three datasets of mammalian nuclei and cytoplasm are used for this work. We show that active deep learning significantly reduces the number of training samples required and also improves the quality of segmentation.

## 1. INTRODUCTION

Cells are the fundamental unit of life. Biologists have sought to explain the core principles over many decades of research. The quest to understand it’s underlying mechanisms continues to this day. A fundamental problem in many studies of cell biology is cell segmentation. Analysis of intracellular processes and understanding of cell physiology is heavily dependent on the quality of the cell segmentation.

Automated cell segmentation has come a long way over the last 50 years [1]. Available solutions draw upon standard techniques – filtering, thresholding, morphological processing and watershed transform from computer vision. Supervised machine learning algorithms have proved to be successful in this space [2–3]. Recent advances in deep learning, particularly Convolutional Neural Networks (CNN) [4], show promise for their application in the field of cell segmentation. CNNs have been shown to be very successful in image classification tasks. They have recently been used for semantic segmentation – the assignment of labels to individual pixels of the image in a computationally efficient manner [5]. It was demonstrated in the 2015 ISBI cell tracking challenge, that CNNs could be used to accurately perform cell segmentation [6]. Another successful attempt was made at performing cell segmentation and subsequent quantitative analysis in live-cell imaging experiments using CNNs [7].

A key component of performing segmentation using supervised learning is annotation. Annotation consists of labeling the regions of interest in the image. This allows the algorithm to learn and quantify the difference between foreground and background. Unfortunately, annotation is a tedious and time-consuming process. Typically, this is done by experts, even though attempts at using crowdsourcing methods are being explored [8]. It is, therefore, important to explore potential ways of reducing the burden of human annotations.

We propose the use of active learning for this purpose. The idea behind active learning is that the learning algorithm will perform better if it is allowed to choose the samples from which it learns. The learner poses queries to the oracle (human or machine) for labeling one or more unlabeled samples. A simple query framework known as uncertainty sampling is used in this work.

We show that using a conjunction of these two methods – CNNs and uncertainty sampling, the amount of training data can be considerably reduced to 1/6^th^ of the original training size, on average. In addition, we also demonstrate that active learning improves the quality of segmentation. Our experiments are performed on 3 datasets consisting of mammalian nuclei and cytoplasm. The paper is organized as follows. Section 2 describes the data that we use for performing the experiments. Section 3 provides a brief explanation of the two methods that are used in this work. Section 4 contains a detailed description of the experiments. Sections 5 and 6 conclude with discussion and future work respectively.

## 2. DATA

Our data sets consist of three mammalian cell lines – NIH-3T3 (mouse embryonic fibroblasts), MCF10A (human breast epithelial cells) and HeLa-S3 (cervix adenocarcinoma). Data and annotations are publicly available, see [7]. The images are in the form of fluorescent microscopic images of nuclei and phase microscopic images of cytoplasm. ImageJ was used for annotating the training dataset and classify each pixel as belonging to the boundary, interior or exterior with respect to the cell. An example of a MCF10A sample is shown in Fig. 1.

**Fig. 1:**
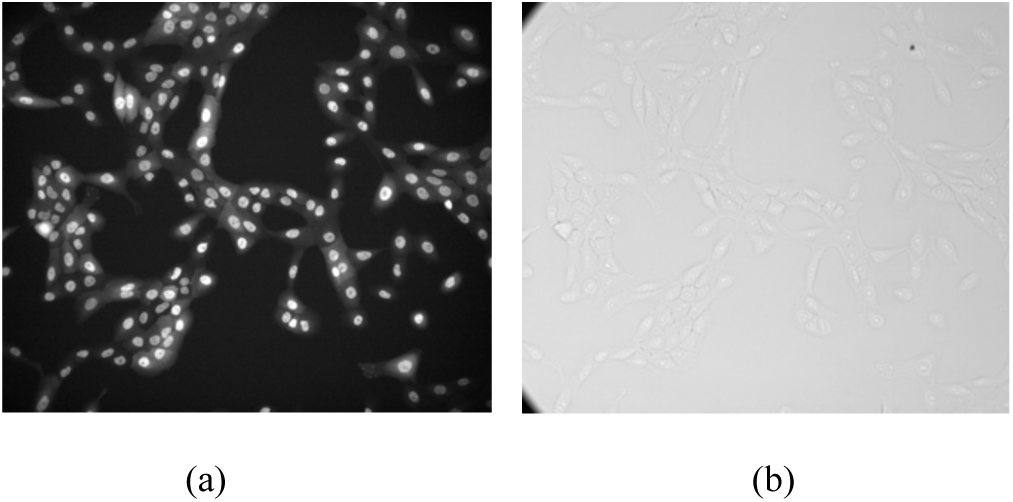
Example of MCF10A cell lines (a) Fluorescent microscopic sample of nuclei (b) Phase microscopic sample of the cytoplasm.

A pixel based classification framework is used to approach the segmentation. The pixels are classified based on the patches around them. The patch-size is set to 61x61 pixels around each pixel. The training data, which are in the form of patches, are augmented using random rotation (0,90, 180, and 270 degrees) and flipping.

## 3. METHODS

Active learning, in particular uncertainty sampling is used is used in conjunction with CNNs. We use two additional methods to fine-tune the segmentation. Active contours [9] is used to fine-tune the segmentation. The methodology in [5] is also leveraged for labeling pixels in computationally efficient manner using d-regularized sparse kernels.

### 3.1. Uncertainty sampling

The main hypothesis in this framework is that the learning algorithm chooses the instances from which learns. It overcomes the labeling bottleneck by asking queries in the form of unlabeled instances to be annotated by an oracle (annotator). It therefore aims at achieving high accuracy using as few labels as possible, thus minimizing the cost of getting labeled data.

Fig. 2 shows a diagrammatic representation of active learning. In uncertainty sampling, the learner queries the user or annotator to label the most informative samples. Three class classification is performed in this work. The algorithm queries the instance whose prediction is the least confident:

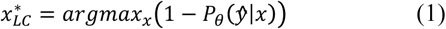

where, 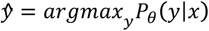 or the class label with the highest posterior probability under the model *θ*. Samples whose highest posterior probability is less than a threshold (0.5) are selected. This method is able to filter and select only those instances that the model is uncertain about predicting.

**Fig. 2:**
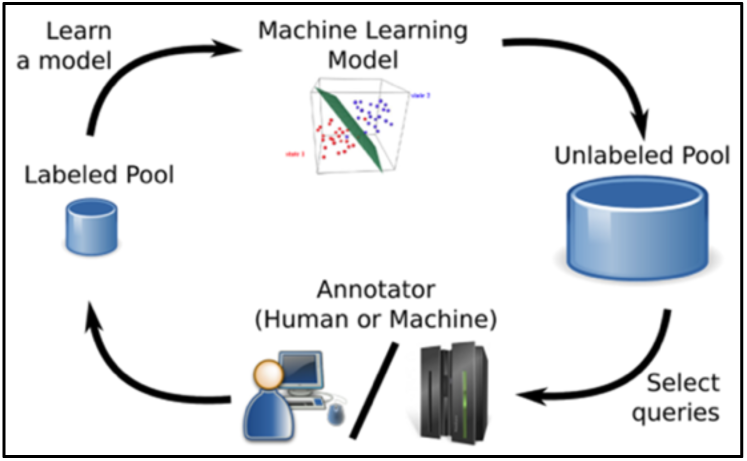
Active learning using uncertainty sampling

### 3.2. Convolutional neural networks

Image segmentation of cells can be converted to an image classification problem. A manually annotated image consists of each pixel being identified as either exterior, boundary or interior pixel. A training dataset may be constructed by sampling a small region around each pixel and assigning the resulting image that pixel’s respective class. The image segmentation task is therefore effectively reduced to finding a classifier that can distinguish between the three classes and can classify new images in a similar manner. Convolutional neural networks is the classifier that does this task. CNN is a deep feed-forward neural network used to analyze visual images. It consists of an input and output layer, as well as multiple hidden layers. The hidden layers consist of convolutional, pooling, non-linear or fully connected layers among others.

The parameters of the model used is the same as that used in [7].

## 4. EXPERIMENTS AND RESULTS

In this section, we describe the experiments and results that were performed in our analysis.

### 4.1. Hyper-parameters

In addition to the parameters of the CNN, there are a number of parameters that control the algorithms themselves. These are known as hyper-parameters. Most of the hyper-parameters are pre-set based on the recommendations from the original papers and common sense. These hyper-parameters are shown in the following table.

**Table 1:** Hyper-parameters of the algorithms.

| Hyper-parameters |  |
| --- | --- |
| Algorithm | Hyper-parameters |
| CNN | Input channels (2), Convolutional (7), batch-normalization (7), rectified linear activation (7), max-pooling (3), fully-connected (2), soft-max activation (1), learning rate (0.01), learning rate decay ($10^{-6}$), number of epochs (25), batch-size (32), momentum (0.9), optimization (stochastic gradient descent) |
| Uncertainty sampling | Probability threshold (0.5) |
| Active contours | Smoothing (1), number of iterations (120) |

The architecture of the CNN is the same as that used in [7]. The input to the CNN is a 2 channel 61x61px sized patch. The two channels represent the nuclei and cytoplasmic patch of the image sample. A CNN is also trained separately on the nuclear patches for segmenting the nuclei. In this case the input is only the nuclear channel. The threshold for uncertainty sampling is set at 0.5. The probability for picking a class at random in a 3-class classification is 0.33. Therefore, if the classifier outputs a probability that is between 0.33 and 0.5, it indicates that it is uncertain about that particular sample. Exploring the space of hyper-parameters using optimization techniques for this task is a possibility for future work.

### 4.2. Algorithm

Pixel based classification is performed for segmentation. Patches around the pixel are sampled from the image and each patch is labeled as either *interior*, *boundary* or *exterior* depending on the location of the pixel with respect to the cells. The classes are balanced by under-sampling. This means the number of patches are determined by the class having the lowest number of samples (this is usually the *boundary* class). The maximum number of samples is set to 1000000. The algorithm for performing the active learning experiment is described in Fig. 3.

**Fig. 3.**
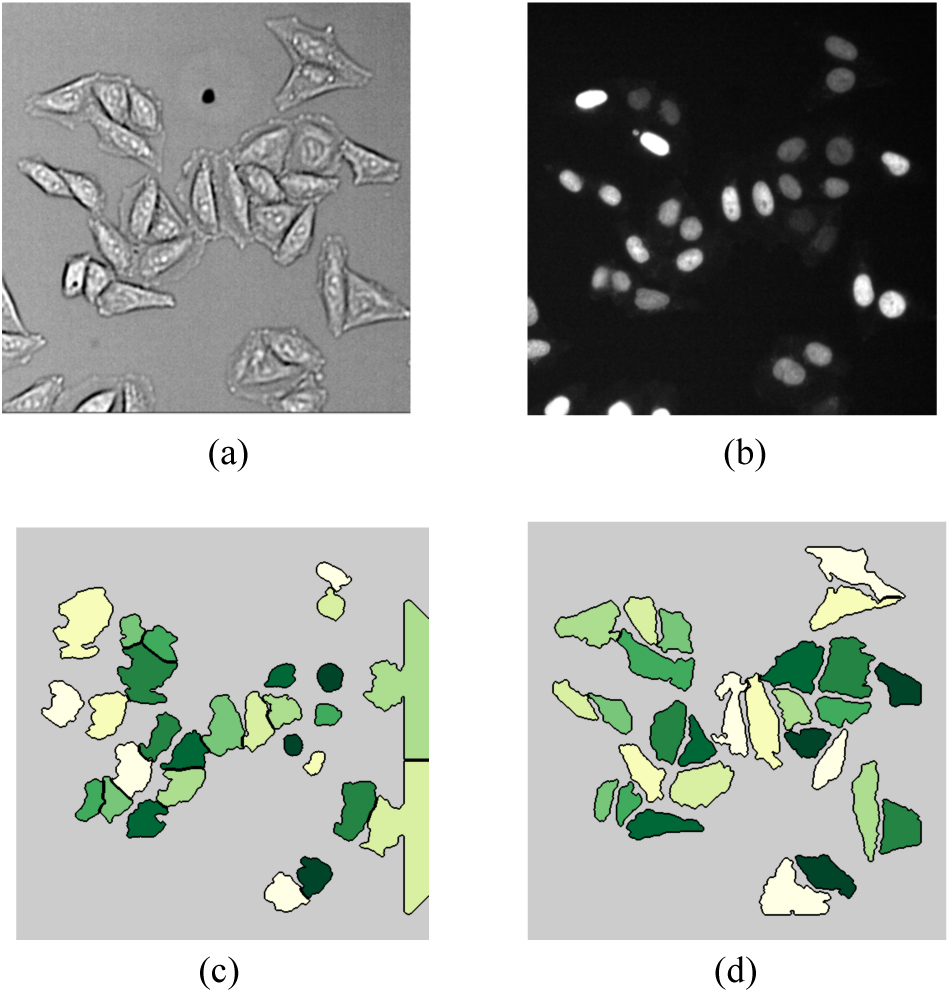
Algorithm for active deep learning

### 4.3. Experimental setup

Two sets of experiments are performed. The first experiment, which we denote as the control experiment simply trains the CNN on 90% of the training data and 10% of the data is held out for testing the classification metrics. The second experiment is the implementation of the algorithm described in the previous section. In addition to the active learning experiment, we also perform a baseline experiment, where uncertainty sampling is not applied after every batch. The results of both these experiments are described in the following section.

### 4.4. Results

Table 2 shows the results for comparing the control experiment and the active learning model obtained after training on the final (9^th^) batch of the algorithm in Fig. 4.

**Table 2.**
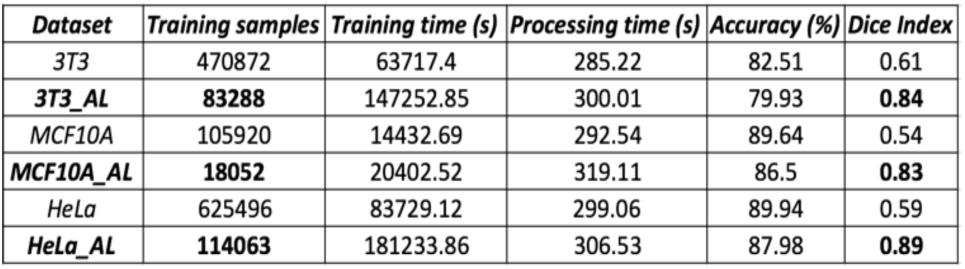
Results

| Dataset | Training samples | Training time (s) | Processing time (s) | Accuracy (%) | Dice Index |
| --- | --- | --- | --- | --- | --- |
| 3T3 | 470872 | 63717.4 | 285.22 | 82.51 | 0.61 |
| 3T3_AL | 83288 | 147252.85 | 300.01 | 79.93 | 0.84 |
| MCF10A | 105920 | 14432.69 | 292.54 | 89.64 | 0.54 |
| MCF10A_AL | 18052 | 20402.52 | 319.11 | 86.5 | 0.83 |
| HeLa | 625496 | 83729.12 | 299.06 | 89.94 | 0.59 |
| HeLa_AL | 114063 | 181233.86 | 306.53 | 87.98 | 0.89 |

**Fig. 4:**
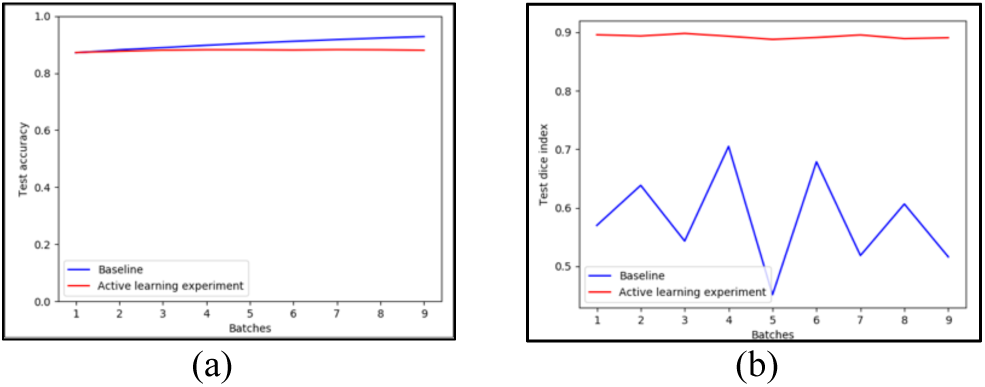
Metrics with respect to number of batches (a) Accuracy (b) Dice index.

From the results in Table 2, we can see that the active learning experiment (denoted by ‘*_AL’) requires much less training data to achieve similar accuracy and better segmentation metrics (dice index). On average, the number of training samples in the active learning experiment are reduced to 1/6^th^ of the number of samples in the control experiment.

It is to be noted that the segmentation metrics are calculated over the entire image and calculated over a single image. Active contours [9] is applied before obtaining the segmentation maps. This might explain the difference in the accuracy and segmentation metrics. This is also observed in the metric plots on the *HeLa* dataset in Fig. 6. The classification metrics with respect to the number of batches are plotted in Fig. 4 (a) comparing the active learning and the baseline experiments. This result is on the 10% held out test data in the algorithm from Fig. 3. Fig. 4(b) shows the segmentation metrics in terms of the dice index on a separate test image. Even though the test is on a single image, active learning does a better job at segmentation than the baseline experiment. It is also more robust than the baseline experiment. This may be explained by the fact that few samples are added to the training of the active learning models on each batch.

The training time for active learning experiments is more than the control. This is because, in the active learning experiment, the models are repeatedly trained at each iteration with the new informative samples. In contrast, in the control experiment, the CNN is trained once for the entire training data. This explains why the training time is more for active learning.

The segmentation maps for an image from the *HeLa* dataset are shown in Fig. 5. Fig. 5(a) and 5(b) are the raw cytoplasm and nuclei images respectively. Fig. 5(c) is the output of the segmentation from the control experiment. Fig. 5(d) is the segmentation map for the model trained by the active learning experiment after the 9^th^ batch of the algorithm in Fig. 3.

**Fig. 5:**
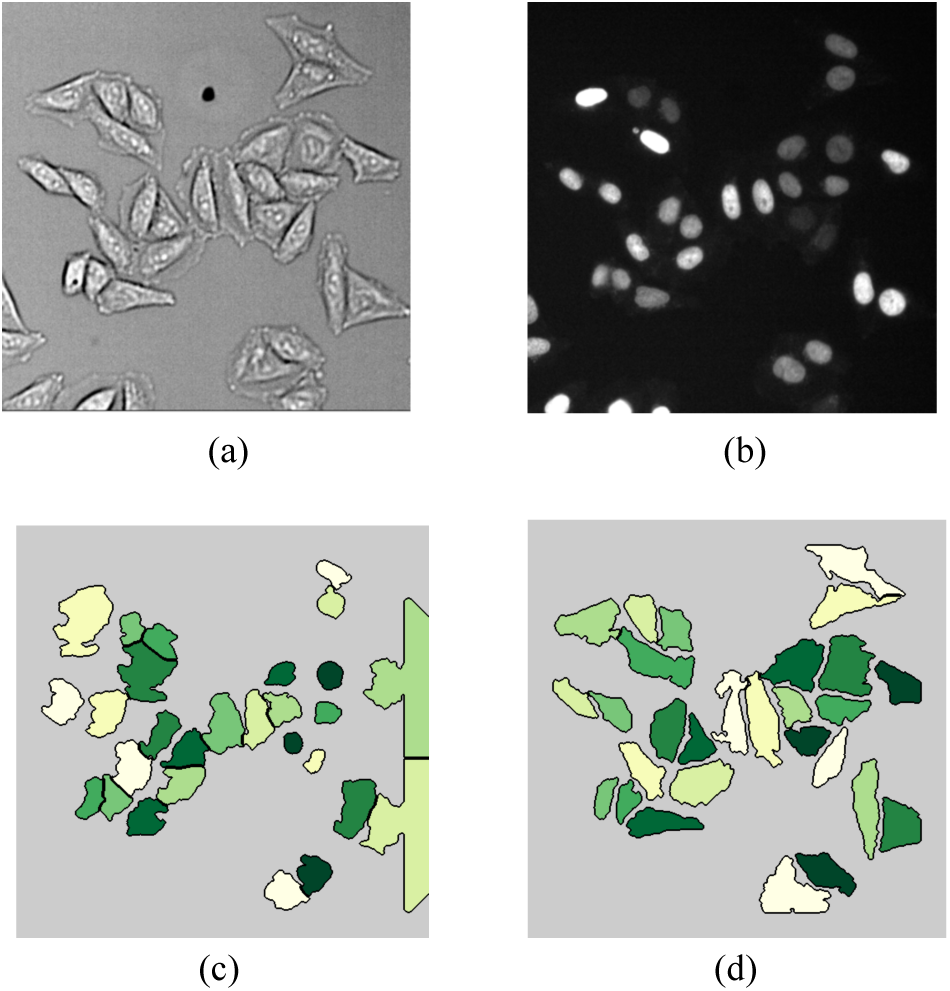
Segmentation outputs. (a) Phase cytoplasmic image (b) Fluorescent nuclei image (c) Segmentation from control experiment. (d) Segmentation from active learning experiment.

We can clearly see that the active learning method does a much better job at segmentation than the control experiment (without active learning).

## 5. CONCLUSION

An active learning based approach may be used to reduce the annotation burden on researchers and medical professionals for segmentation tasks. Very few training samples are required for achieving good quality segmentations. From the results, we can see that the number of training samples may be reduced to 1/6^th^ of the original training size on average. Also, since the training is done on informative samples determined by the active learning algorithm, the quality of the segmentation also improves significantly.

## 6. FUTURE WORK AND ACKNOWLEDGEMENTS

There are a number of directions to be pursued in terms of future work. The first is testing the methods on a new and harder dataset and domain. This is because the methods described in the paper maybe easily generalizable to other domains and datasets. The second is using unsupervised learning to warm-start the active learning process. The first batch of training in the algorithm described in section 4.2 may be sampled in a more intelligent manner using clustering. This work was successfully completed at IBM Research, Almaden.

SKB and SB thankfully acknowledge the National Science Foundation for financial support under Grant No. DBI-1548297. Any opinions, findings and conclusions or recommendations expressed in this material are those of the author(s) and do not necessarily reflect the views of the National Science Foundation.

## REFERENCES

[1] E. Meijering, "Cell segmentation: 50 years down the road [life sciences]," IEEE Signal Processing Magazine vol. 29, no. 5, pp. 140–145, Sep 2012.

[2] K.Z. Mao, P. Zhao, P.H. Tan, “Supervised learning-based cell image segmentation for p53 immunohistochemistry,” IEEE Transactions on Biomedical Engineering, vol. 53 no. 6, pp. 1153–1163, Jun 2006.

[3] L. Bertelli, T. Yu, D. Vu, B. Gokturk, “Kernelized structural SVM learning for supervised object segmentation,” IEEE Conference on Computer Vision and Pattern Recognition, pp. 2153–2160, Jun 2011.

[4] A. Krizhevsky, I. Sutskever, and G. E. Hinton, "Imagenet classification with deep convolutional neural networks," Advances in neural information processing systems, pp. 1097–1105, 2012.

[5] H. Li, R. Zhao, X. Wang, “Highly efficient forward and backward propagation of convolutional neural networks for pixelwise classification,” arXiv preprint vol. 1412 no. 4526, Dec 2014.

[6] O. Ronneberger, P. Fischer, T. Brox, “U-Net: Convolutional Networks for Biomedical Image Segmentation,” Medical Image Computing and Computer-Assisted Intervention, pp. 234–41, 2015.

[7] D.A. Van Valen, T. Kudo, K.M. Lane, D.M. Macklin, "Deep Learning Automates the Quantitative Analysis of Individual Cells in Live-Cell Imaging Experiments," PLoS Computational Biology, vol. 12 no. 11, Nov 2016.

[8] A. Hughes, J.D. Mornin, S.K. Biswas, D.P. Bauer, Bianco, S. and Z.J. Gartner, “Quantius: Generic, high-fidelity human annotation of scientific images at 105-clicks-per-hour,” bioRxiv, p. 164087, 2017.

[9] P. Marquez-Neila, L. Baumela, L. Alvarez, “A morphological approach to curvature-based evolution of curves and surfaces,” IEEE Transactions on Pattern Analysis and Machine Intelligence, vol. 36 no. 1, pp.2–17, 2014.

